# scAdapt: Virtual adversarial domain adaptation network for single cell RNA-seq data classification across platforms and species

**DOI:** 10.1101/2021.01.18.427083

**Authors:** Xiang Zhou, Hua Chai, Yuansong Zeng, Huiying Zhao, Ching-Hsing Luo, Yuedong Yang

**Affiliations:** School of Data and Computer Science, Sun Yat-sen University, Guangzhou 510000, China; Sun Yat-sen Memorial Hospital, Sun Yat-sen University, Guangzhou 510000, China; Key Laboratory of Machine Intelligence and Advanced Computing (Sun Yat-sen University), Ministry of Education, China

## Abstract

**Motivation:** In single cell analyses, cell types are conventionally identified based on known marker gene expressions. Such approaches are time-consuming and irreproducible. Therefore, many new supervised methods have been developed to identify cell types for target datasets using the rapid accumulation of public datasets. However, these approaches are sensitive to batch effects or biological variations since the data distributions are different in cross-platforms or species predictions.

**Results:** We developed scAdapt, a virtual adversarial domain adaptation network to transfer cell labels between datasets with batch effects. scAdapt used both the labeled source and unlabeled target data to train an enhanced classifier, and aligned the labeled source centroid and pseudo-labeled target centroid to generate a joint embedding. We demonstrate that scAdapt outperforms existing methods for classification in simulated, cross-platforms, cross-species, and spatial transcriptomic datasets. Further quantitative evaluations and visualizations for the aligned embeddings confirm the superiority in cell mixing and preserving discriminative cluster structure present in the original datasets.

**Availability:** https://github.com/zhoux85/scAdapt.

## 1 Introduction

Single-cell RNA-seq technologies have been successfully employed to generate high resolution cell atlas and to improve our understanding of cellular heterogeneity in human diseases. One major step of single-cell RNA sequencing (scRNA-seq) analyses is cell type identification (Lahnemann, et al., 2020). Typically, cells are first grouped into different clusters, and each cell cluster will be manually assigned to one label based on the uniquely high expression levels of canonical makers. Nevertheless, visual inspection of cluster-specific gene is labor intensive in practice and irreproducible, and the assignments of cell types require expert knowledge of canonical makers (Luecken and Theis, 2019). Thus, it’s necessary to develop automated computational methods for cell annotations.

A growing list of classification methods have been developed to annotate cells based on public data of known cell types (Abdelaal, et al., 2019). The most typical methods are similarity-based methods that assign cell labels through scanning reference cell databases for similar cells. For example, SingleR (Aran, et al., 2019), and CHETAH (de Kanter, et al., 2019) used Spearman correlation for similarity measurement. The scmap (Kiselev, et al., 2018) combines three metrics, cosine distance, Pearson correlation, and Spearman correlation, to quantify the closeness between query cell and the centroid of each reference cell cluster. Though these methods are robust, they cannot reflect the complex non-linear relations between genes. To overcome this issue, the machine-learning based methods are proposed to train models on the reference dataset and to use the trained models for predicting cell labels. For example, scPred (Alquicira-Hernandez, et al., 2019) takes advantage of singular value decomposition to obtain small number of informative features and uses the features to train a support vector machine (SVM) model. In singleCellNet (Tan and Cahan, 2019), the reference data are pair-transformed into binary matrix that is then used to train a Random Forest classifier. Seurat (Stuart, et al., 2019) identifies anchoring cell pairs by projecting query cells onto precomputed reference principal component analysis (PCA) structure and used these anchors to train a weighted vote classifier for cell annotation. However, machine learning techniques applicable to test sets following the same distribution as the training set (Ganin and Lempitsky, 2015), and thus do not always work well on single cell data due to batch effects or biological factors (e.g. treatments, individuals, species) difference between datasets.

To solve the distribution mismatch between samples, many methods have been proposed to align cell distribution, such as fastMNN (Haghverdi, et al., 2018), Harmony (Korsunsky, et al., 2019), LIGER (Welch, et al., 2019). However, most of these methods do not support label prediction, and other cell annotation tools such as singleCellNet and SingleR running on the aligned data didn’t show much improvement due to the transformation of gene expression according to previous benchmark analyses (Abdelaal, et al., 2019; Huang, et al., 2020). Seurat is the only method to support a joint batch effect removal and cell annotation. When the batch difference is obvious, Seurat has the option to learn an aligned subspace across datasets using canonical correlation analysis (CCA) instead of PCA, where anchors are identified for classification. Although taking batch difference into consideration and showing improvement in practice, Seurat suffers from two limitations. First, the integration is unsupervised without effectively using the cell-type information in reference data for aligning cell clusters, and it may mismatch cells of different cell types across datasets. Second, since distribution-alignment and label projection are optimized independently, the features for sample alignment are not optimal for cell classification. Therefore, it should be beneficial to combine sample alignment and cell annotation in one step.

In computer vision community, domain alignment and classifier training can be joint performed through domain adaptation network that has been shown to enhance the generalization of the classification model (Wang and Deng, 2018). A typical domain adaptation framework is to reduce the distribution mismatch of the latent feature via domain adversarial learning (Ganin and Lempitsky, 2015). It can also be readily applied to the scenario of cross-batch single cell annotation, where scRNA-seq data from different batches are considered as different domains (Ge, et al., 2020). On the other hand, semi-supervised learning (SSL), can leverage additional information from the unlabeled data to better estimate the decision boundary between the different classes, and thus improves the classifier’s accuracy (Ouali, et al., 2020). SSL can also be used to alleviate the adverse impact of domain discrepancy by training the classifier on labeled source and unlabeled target data jointly (Cui, et al., 2020). Virtual adversarial training (VAT) has been widely used in many SSL tasks and achieves state-of-the-art performance (Miyato, et al., 2018). It can enhance classifier’s robustness with respect to random and local perturbations or noises in the inputs.

Combining domain adaptation and VAT-based semi-supervised learning, we developed scAdapt, to make use of both labeled source and unlabeled target data for improving classification performance. Here, our domain adaptation network includes not only the adversary-based global distribution alignment, but also category-level alignment (Xie, et al., 2018) to preserve the discriminative structures of cell clusters in low dimensional feature (i.e., embedding) space. We demonstrate that scAdapt compares favorably to existing methods for classification and batch correction in simulated, cross-platforms, cross-species, and spatial transcriptomic datasets. Further quantitative evaluations and visualizations for the aligned embeddings confirm the superiority in cell mixing and preserving discriminative cluster structure present in the original datasets.

## 2 Methods

### 2.1 Datasets and preprocessing

#### Simulated data

We used the *R* package “*Splatter”* (Zappia, et al., 2017) to generate simulated scRNA-seq counts data of different batches with similar cell type compositions. We simulated two batches with 2000 and 1000 cells considered as source and target dataset, respectively, and each cell has 10000 genes. Each batch was uniformly split into four cell groups with cell proportion set to 0.25 by the parameter *group.prob*. To simulate datasets with different magnitudes of batch effects, we adjusted the batch parameter *batch.facLoc* and *batch.facScale* with increasing values [0.2, 0.4, 0.6, 0.8, 1.0] where larger values corresponding to stronger batch effects. For brevity, we set *batch.facLoc* = *batch.facScale*. To simulate datasets with different magnitudes of clustering difficulty, we set the parameter de.fracScale to 0.2 for simulated datasets with weak clustering signal and 0.3 for simulated datasets with strong clustering signal. Simulation were run five times with different random seeds and average results were reported. For other parameters, default values were used unless otherwise specified.

#### Cross-platforms datasets

The human Peripheral Blood Mononuclear Cells (PBMC) scRNA-seq data were retrieved directly from the SeuratData package with dataset name “pbmcsca” (Ding, et al., 2019). The data consists of seven batches from seven different sequencing platforms. We removed the cells annotated as “Unknown” and the resulting datasets contains a total of 30975 cells and each cell has 33694 genes. We combined the data from the 10x Chromium (v2) and 10x Chromium (v3) platform as source data and the rest five platforms: CEL-Seq2 (CL), Drop-seq (DR), inDrop (iD), Smart-seq2 (SM2), Seq-Well (SW) as target data. As a result, we have five pairs of cross-platform datasets: 10x-CL, 10x-DR, 10x-iD, 10x-SM2, 10x-SW. For all the datasets, raw counts were extracted from the Seurat object for further processing.

#### Cross-species datasets

The human and mouse pancreas data were downloaded from SingleCellNet GitHub page where five ready-to-use datasets are provided. For data batch generated by Baron, Segerstolpe and Tabula Muris cell atlas, raw counts are provided for further processing. For datasets from Murano and Xin, normcounts are provided. Following the filtering step in previous benchmark study (Tran, et al., 2020), we removed the cells labeled as “unclear”, “co-expression”, “unclassified”, “unclassified endocrine”, “alpha.contaminated”, “beta.contaminated”, “delta.contaminated” or “gamma.contaminated”. “activated_stellate,” “PSC”, and “quiescent_stellate” cells were merged into “stellate”. The resulting datasets contain a total of 17,574 cells. To obtain compatible gene names for cross-species analysis, we used the homologous genes provided by SingleCellNet to convert gene names and only the intersection gene set between the human data and mouse data were kept. To construct a large source with enough training samples and cover more cell types in the source, we combined the mouse data from the Baron and Tabula Muris as source data.

#### Spatial transcriptomic datasets

We downloaded two mouse brain (hypothalamic preoptic region) datasets from Gene Expression Omnibus (GSE113576) and Dryad repositories, respectively (Moffitt, et al., 2018). The spatial transcriptomic dataset has 64,373 cells measured with spatially resolved multiplexed error robust fluorescence in situ hybridization (MERFISH) and the scRNA-seq dataset has 30,370 cells measured by 10x Chromium. 10x data has full transcriptome with 22,067 genes, while MERFISH data has only 154 targeted genes. We combined the two datasets with the 154 intersecting genes.

#### Preprocessing

Seurat R package (version 3.2.0) was used for preprocessing. For both the simulated and real datasets (except for Murano, Xin, and MERFISH where counts are already normalized), the counts matrix were normalized by the NormalizeData function in Seurat with default ‘LogNormalize’ normalization method and a scale factor of 10,000. Top 2000 highly variable genes were selected based on the log-normalized counts using the FindVariableFeatures function with default ‘vst’ method. For real datasets, the cell-type annotations from the corresponding publications were considered as the ground truth for evaluations. Because the brain data has pre-selected markers, we did not select variable gene, but used all the 154 intersecting genes.

The datasets analyzed in this study are summarized in Table S1.

### 2.2 The architecture of scAdapt

Our scAdapt model includes two modules. Classification module, based on cross-entropy loss and virtual adversarial training loss, aims to improve the accuracy of cell annotation using both labeled source and unlabeled target data. Batch correction module contains two loss. The adversarial domain adaptation loss aims to reduce distribution discrepancy at embedding space of source and target, while the semantic alignment loss can make the embeddings better clustered and more separable. We optimized these two modules jointly in order to improve domain alignment and final classification simultaneously.

The overall structure of scAdapt is illustrated in Fig. 1. It consists of a feature extractor G with two hidden layers, a domain classifier D with two hidden layers, and a label predictor F with a linear output layer followed by a softmax operation. The input includes source gene expression matrix 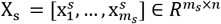 of *m*_*s*_ labeled cells with 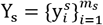 being the corresponding labels and target gene expression matrix 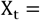 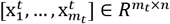 of *m*_*t*_ unlabeled cells, where *n* is the number of common genes shared by the source and target data. In domain adaptation setting, X_s_ and X_t_ are assumed to be different but related (Wang and Deng, 2018).

**Fig 1.**
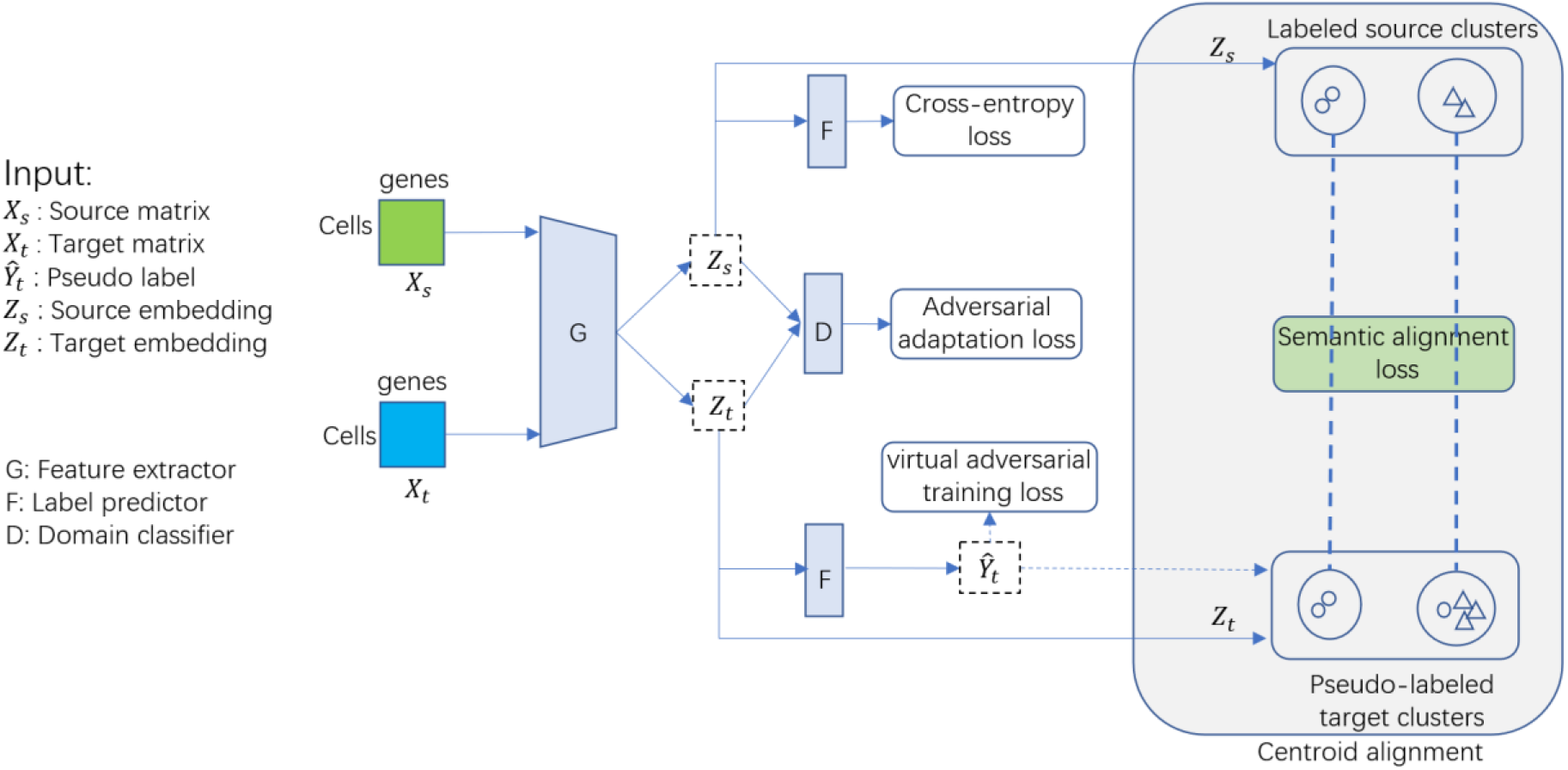
The overall structure of scAdapt for cell type classification and batch correction on source and target dataset. With source and target data as input, the feature extractor G learns to capture low-dimensional embedding *Z*_*s*_ and *Z*_*t*_ which are then used to train the label predictor F with the cross-entropy loss and virtual adversarial training loss, respectively. At the embedding space, batch correction is achieved at global- and class-level: adversarial domain adaptation loss is employed to perform global distribution alignment and semantic alignment loss minimizes the distance between the labeled source centroid and pseudo-labeled target centroid. The target pseudo label 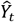 is estimated by label predictor F.

To minimize the source sample classification error with known labels, standard cross-entropy loss is used as below:

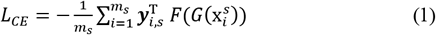

where ***y***_*i,s*_ ∈ *R*^*K*×1^ is one-hot encoded vector of 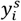 and *K* is the number of class.

We use virtual adversarial training (VAT) to incorporate the information of data distribution from unlabeled data, which can better estimate the decision boundary between different classes (Ouali, et al., 2020). VAT is an effective data augmentation technique which do not need prior label information and is hence applicable to semi-supervised learning. It assigns similar labels to each input data and its neighbors in the adversarial direction where the perturbation will alter the model’s output distribution the most. Then the model is robust to small perturbations or noises in the inputs. The loss function of VAT is given by

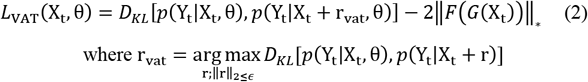

where r_vat_ denotes the virtual adversarial perturbation maximizing the difference between the model output of perturbed input and non-perturbed input, θ is the model parameter to train. The output distribution is parameterized as *p*(Y_t_|X_t_, θ), and *D*_*KL*_[∙,∙] is KullbackLeibler divergence that measures the difference between two probability distributions. The last penalty term ‖*F*(*G*(X_t_))‖_*_ in (2) is designed to improve both the prediction discriminability and diversity, and ‖∙‖_*_ is the nuclear-norm (Cui, et al., 2020).

To learn the domain-invariant features, adversarial adaptation loss is adopted, where the feature extractor G and domain classifier D are trained by playing a two-player minimax game: the first player is domain classifier which distinguishes whether the feature is from the source domain or target domain, and the second player is feature extractor which aims to output domain-invariant features to confuse the domain classifier. Domain alignment is expected when the game reaches an equilibrium. Formally, the domain classifier D is trained by minimizing the binary cross-entropy loss

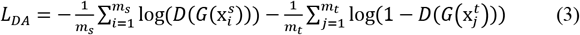

while the feature extractor G is trained to maximize the *L*_*adv*_ loss (fool the domain classifier D). In order to update the parameters of D and G simultaneously, gradient reverse is used to flip the sign of the gradient between D and G during backpropagation (Ganin and Lempitsky, 2015).

Besides global domain-invariance, discriminability must also be preserved, which ensures the embeddings of same class but different domains are mapped nearby. An intuitive solution is to perform semantic alignment for samples of each class directly. However, explicit alignment for each class is impossible since no label information provided for target domain. We approach the problem by assigning pseudo labels to target samples with the classifier F and then explicitly align the centroid for each class in source and target domain (Xie, et al., 2018). The centroid is defined as the mean embedding of each class. For each target class, all samples with correct or wrong pseudo labels are used for centroid calculation, and thus the noise or bias brought by partial false pseudo labels are expected to be suppressed by correct pseudo labels with a dominating portion. We formulate the following semantic alignment loss to minimize the distance between the target centroids and their corresponding source centroids:

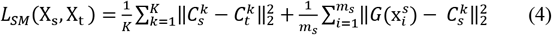

where 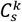 and 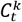 denote the source and target centroids. The second term of (4) is designed to penalize big intra-class distances and enforce better cluster compactness (Wen, et al., 2016).

The overall loss function can be formulated as:

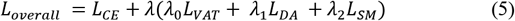

where *λ*_0_, *λ*_1_, and *λ*_2_ are the regularization coefficients controlling the contribution of virtual adversarial training, global domain alignment, and semantic alignment to the total loss function, respectively.

### 2.3 Identifying cell-type-important genes

Since neural networks are often considered as black box models with no clear interpretation, examining the importance of each gene relative to classification output is favorable for understanding the reason behind classification decisions. We identified key genes for each cell type by activation maximization method (Simonyan, et al., 2013). Formally, let *θ* be the fixed model parameters after training the network, and *h*_*i*_(*θ*, *x*) be the activation of *i*-th neuron in the last layer of neural network with input *x*, i.e., the classification score for cell type *i*. Activation maximization looks for input patterns which maximize the classification score:

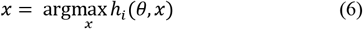

A locally-optimal solution of (6) can be found through gradient ascent in the input space, where the gradient of *h*_*i*_(*θ*, *x*) with respect to *x* are computed to iteratively update the input *x*. It should be noted that the optimization is performed with respect to the input *x*, which is different from the training procedure of neural network for optimizing the model parameters *θ*. The input *x* was initialized with a zero vector and updated for 100 iterations with learning rate set to 1. The changes of resulting *x*\* compared with the initialization values were calculated as the gene importance score. To evaluate whether the identified top-important genes are reliable, we selected the top genes with the largest importance score for each cell type and compared them with cell-type markers in the PanglaoDB database (Franzén, et al., 2019) and the marker gene reported in original publication (Moffitt, et al., 2018). We also performed Gene Ontology (GO) enrichment analysis on these selected genes, using the R package clusterProfiler (Yu, et al., 2012).

### 2.4 Hyper-parameters setting

All the neural network layers are fully-connected. The two hidden layers of feature extractor G have 512 and 256 nodes, respectively. For spatial transcriptomic data with only 154 genes, we set the nodes in each hidden layer as 128 and 128. The size of hidden layers in domain classifier D is set to 1024. Rectified linear unit (ReLU) function is used as activation function for the hidden layers while softmax activation function and sigmoid function applied to the last layer of F and D, respectively. The network is trained by mini-batch stochastic gradient descent with a momentum of 0.9 and weight decay of 5 × 10^−4^. We follow the same annealing strategy of learning rate as described in (Ganin and Lempitsky, 2015), i.e. the learning rate *η*_*p*_ is adjusted following *η*_*p*_ = *η*_0_/(1 + *ap*)^*b*^, where *p* is the training progress linearly increasing from 0 to 1, *η*_0_ is the initial learning rate set to 0.001, a = 10, and b = 0.75. The batch size is set to 256. Throughout all experiments, we set regularization parameter *λ*_0_ = 0.1*λ*, *λ*_1_ = *λ*, and *λ*_2_ = *λ*, where the penalty parameter *λ* is updated from 0 to 1 by a progressive schedule according to Ganin et al. (Ganin and Lempitsky, 2015):

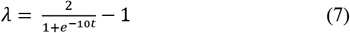

where *t* is the training progress linearly increasing from 0 to 1. With this schedule, the model can first focus on training the model with labeled source data, and then focus on the optimization of virtual adversarial training, global domain alignment, and semantic alignment whose information are noisy and inaccurate at early training stages. The method was implemented based on PyTorch (Paszke, et al., 2019). An open-source implementation of the scAdapt algorithm can be downloaded at https://github.com/zhoux85/scAdapt.

### 2.5 Benchmarking classification methods

To evaluate the performance of scAdapt, we benchmarked it against other cell type annotation tools, including: Seurat V3, scmap, scPred, CHETAH, SingleR, and singleCellNet. For Seurat V3, we used both the CCA-based and PCA-based label propagation to evaluate whether the classification can benefit from aligned data. The default hyperparameters recommended in these annotation tools and accompanying tutorials were used for performance evaluation.

#### Evaluation metric

We evaluated the classification performance of each method using the accuracy score, which is defined as the proportion of correctly annotated cells. We computed the accuracy for each class in the test data and reported averaged accuracy across all the classes. Throughout the evaluation, the previously published cell type annotations provided by original datasets were considered as ground truth.

### 2.6 Benchmarking batch correction methods

Seurat V3, fastMNN, Harmony, and LIGER are used as competing methods. All the tools were run with their default parameters. We evaluated the performance by quantitative measure and visual inspection. Silhouette score and divergence score are used to measure the quality of batch correction (Wang, et al., 2019). An accurate batch correction method should result in a high silhouette score (preserving the original structure of the data) and low divergence score (keeping the same-type cells across batches well mixed). Uniform manifold approximation and projection (UMAP) was used for visualizing cells in a two-dimensional space (Becht, et al., 2018). During the benchmark, all competing methods were run with their default hyperparameters, or the hyperparameters provided in the accompanying tutorials.

#### Evaluation metric

We used divergence score to quantify how well the same population between different batches are mixed after batch correction. A smaller divergence score means better mixing of the same cell population. The quality of mixing is estimated by the universal k-nearest-neighbor (kNN) divergence (Wang, et al., 2009). The kNN divergence between UMAP embeddings *Z*_*s*,*l*_ and *Z*_*t*,*l*_ of class *l* can be formulated as

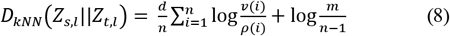

where *d* is dimension size of embeddings, m and n are the number of source and target samples in class *l*, respectively, *ρ*(*i*) is the Euclidean distance between sample *i* and its kNNs in the same batch, and *v*(*i*) is the distance from sample *i* to its kNNs in the other batch. The average kNN divergence over all classes is calculated as divergence score. In all experiments for batch-correction evaluation, *k* was set to 30 for k-nearest-neighbor computation.

Evaluation only by divergence score is not sufficient, since we can obtain a perfect score by randomly mixing the data regardless of the cell type. Thus, we use silhouette score to quantify how well different cell types are separated after batch correction and ensure that datasets integration can conserve true biological signals in original datasets. Let *a*(*i*) denote the average distance between cell *i* to all other cells in the same cluster and *b*(*i*) be the average distance between the cell *i* and cells in the next closest cluster. The distance is calculated using Euclidean distance based the UMAP embeddings of the batch-corrected data. The silhouette coefficient of cell *i* can be formulated as

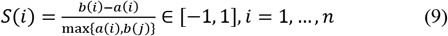

The average silhouette coefficient across all cells can be calculated as silhouette score. A higher silhouette score indicates better cell type assignment.

## 3 Results

To showcase the strength of scAdapt, we analyzed multiple scRNA-seq datasets from simulation tool “*Splatter*”, different species and sequencing platforms, and spatial transcriptomic. The performance of scAdapt was compared with seven cell type classification methods, and four batch correction methods. Our results show that scAdapt consistently outperforms these existing methods in cell type annotation and batch correction.

### 3.1 Performance on simulated dataset

We first evaluated the classification accuracy under different degree of batch effects with weak clustering signal strength (*de.facScale*=0.2) in Fig 2a. We can see that the accuracy of all classification methods decreases with increasing *batch.fracScale*, confirming our speculation that batch effects makes classification challenging. In particular, the accuracy of competing methods drop dramatically as the *batch.fracScale* increases, but the decreasing of Seurat-CCA is not so pronounced at high *batch.fracScale* values. Compared to Seurat-PCA, Seurat-CCA achieved a 78% improvement of accuracy (from 0.39 to 0.69) at the largest *batch.fracScale,* which confirms that batch-correction can enhance the classification model when the batch difference is obvious. Although singleR achieves the second highest performance when *batch.fracScale* <= 0.6, its accuracy demonstrates a sharp drop from 0.85 to 0.61 as *batch.fracScale* changed from 0.6 to 1.0. By comparison, scAdapt always outperforms the competing methods across all batch effects settings and is particularly prominent for large batch difference. The accuracy of scAdapt is perfect (accuracy ≈ 0.98) until the *batch.fracScale* value increased to 0.6, but even then remains high at *batch.fracScale* = 1.0 (accuracy > 0.94). The superior performances show that scAdapt can effectively reduce performance degradation brought by batch difference.

**Fig 2.**
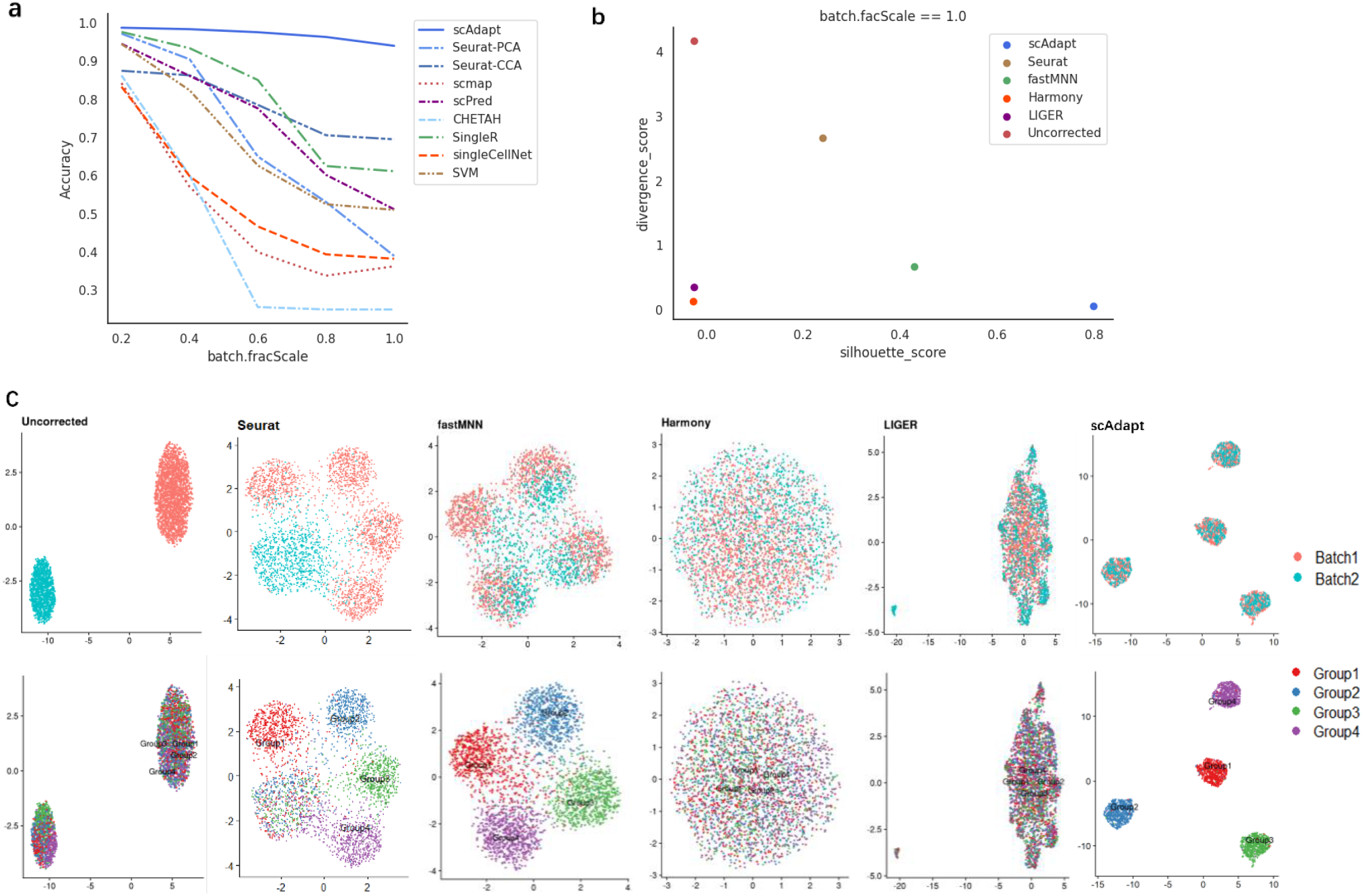
Benchmarking of scAdapt against seven classification methods and four batch correction methods on simulated data with two batches and four different cell types. **a.** Average accuracy under increasing *batch.fracScale* values where larger values corresponding to stronger batch effects. **b.** The integration quality measured by divergence score versus silhouette score at *batch.fracScale* =1.0. Specifically, a lower divergence score means better cell mixing across datasets and a higher silhouette score indicates better cell type assignment. **c.** UMAP plots colored by batch and cell type at *batch.fracScale* =1.0.

Next, we evaluated the performance of scAdapt and other four batch correction methods using the most challenging simulation setting with *batch.fracScale*=1.0 (Fig 2b, c). Ideally, there should be four distinct cell groups (each representing a cell type) in the UMAP visualization after batch correction, and the cells from both batches are well mixed in each group. Visualization of the uncorrected data shows that the cells are grouped by batch distinctly, resulting in lowest silhouette score (−0.026) and highest divergence score (4.18). After removing batch effects by scAdapt, the batch distinctions are effectively removed with the same-type cells across batches uniformly mixed while maintaining the cell type structure in original batches. scAdapt achieves not only the highest silhouette score (0.92) but also presents lowest divergence score (0.01) over others. Although Harmony and LIGER also have low divergence score (≈ 0.10), their silhouette scores are much lower (≈ 0) due to over-correction problem with all cell types mixed together. fastMNN, although produced a proper balance of batch mixing and cell type mixing with a divergence score of 0.75 and silhouette score of 0.43, suffered from under-correction where the cell types across batches are not well aligned despite relatively clear separation between cell types. Seurat produced the highest divergence score (2.5) and low silhouette score (0.24) since it failed to perform cell type alignment and the cell types in Batch 2 are far less discernable. These results suggest improved batch correction by scAdapt, compared with the unsupervised batch correction methods ignoring label information of source data which provide a prior regarding cell-type composition.

To demonstrate the separate contributions of different components in scAdapt, we performed ablation study with *batch.fracScale*=1.0 by evaluating three variants of scAdapt: Baseline refers to scAdapt without VAT and domain adaptation (DA), which means the model was only trained on the source batch; Baseline+VAT and Baseline+DA refer to Baseline including DA and VAT, respectively. From Fig S1a, we can see that the inclusion of VAT can improve the classification notably, and DA is beneficial to batch correction, compared with Baseline. Combing these two components can further enhance the performance of batch correction through guiding cell type alignment with more accurate pseudo labels. We also visualized the embeddings of scAdapt and its three variations by UMAP in Fig S1b. In Baseline, the two batches are completely mismatched and the four clusters in target batch stay too close to each other, which makes the classification of target examples hard. Using the auxiliary information from target, Baseline+VAT makes the cell types discriminated well but ignores alignment across batches. Baseline+DA aligns the cell types correctly, but the group 1 and group 2 are not well separated. For Baseline+DA+VAT, the same cell types across batches are perfectly aligned while different cell types are well distinguished, confirming the necessity of combining VAT and DA for batch correction. We also evaluated the performance on datasets with strong clustering signal strength (*de.facScale*=0.3) where the uncorrected data have low cell type noise and demonstrate distinguishable cluster structure in each batch (Fig S1c). As expected, all classification methods and batch correction methods (except LIGER) show improved performance and scAdapt still outperforms the competing methods.

### 3.2 Performance on cross-platform datasets

In realistic scenario, the source and target datasets are often generated from different experimental platforms by different labs. To evaluate the performance of scAdapt on this realistic setting, we conducted cross-platform test on five paired source-target PBMC datasets where we mapped the cell types from source dataset to target dataset. In this setting, each dataset is profiled by different sequencing platforms.

Fig 3a shows the accuracy score on the five pairs of source-target datasets. scAdapt shows a slight edge over Seurat-CCA by average accuracy, and consistently outperforms the other seven methods across the five test pairs, indicating that integrating source and target dataset can make the classification method resilient against batch effects. The heatmap of confusion matrices of classification results show that scAdapt has a more balanced performance on each cell type with minimum accuracy > 0.74, compared to other methods with minimum accuracy ranging from 0.31 to 0.62 (Fig S2a). The performance drop of competing methods mainly come from the misclassification of closely related cell types. For example, the second ranked Seurat-CCA method incorrectly assigned 21% of Cytotoxic T cell as CD4+ T cell, and 10% of them to Natural killer cell. For completeness, we also ran CHETAH, SingleR, singleCellNet, and SVM with gene expression data corrected by Seurat V3 (other methods failed to run over corrected datasets) and observed that using Seurat V3 correction can improve the performance of CHETAH, SingleR, singleCellNet, and SVM by 15%, 4%, 3%, 4%, respectively (Fig S2b). It’s worth noting that the improved accuracies of CHETAH (0.83), SingleR (0.84), singleCellNet (0.8), and SVM (0.8) are still lower than that of scAdapt (0.86).

**Fig 3.**
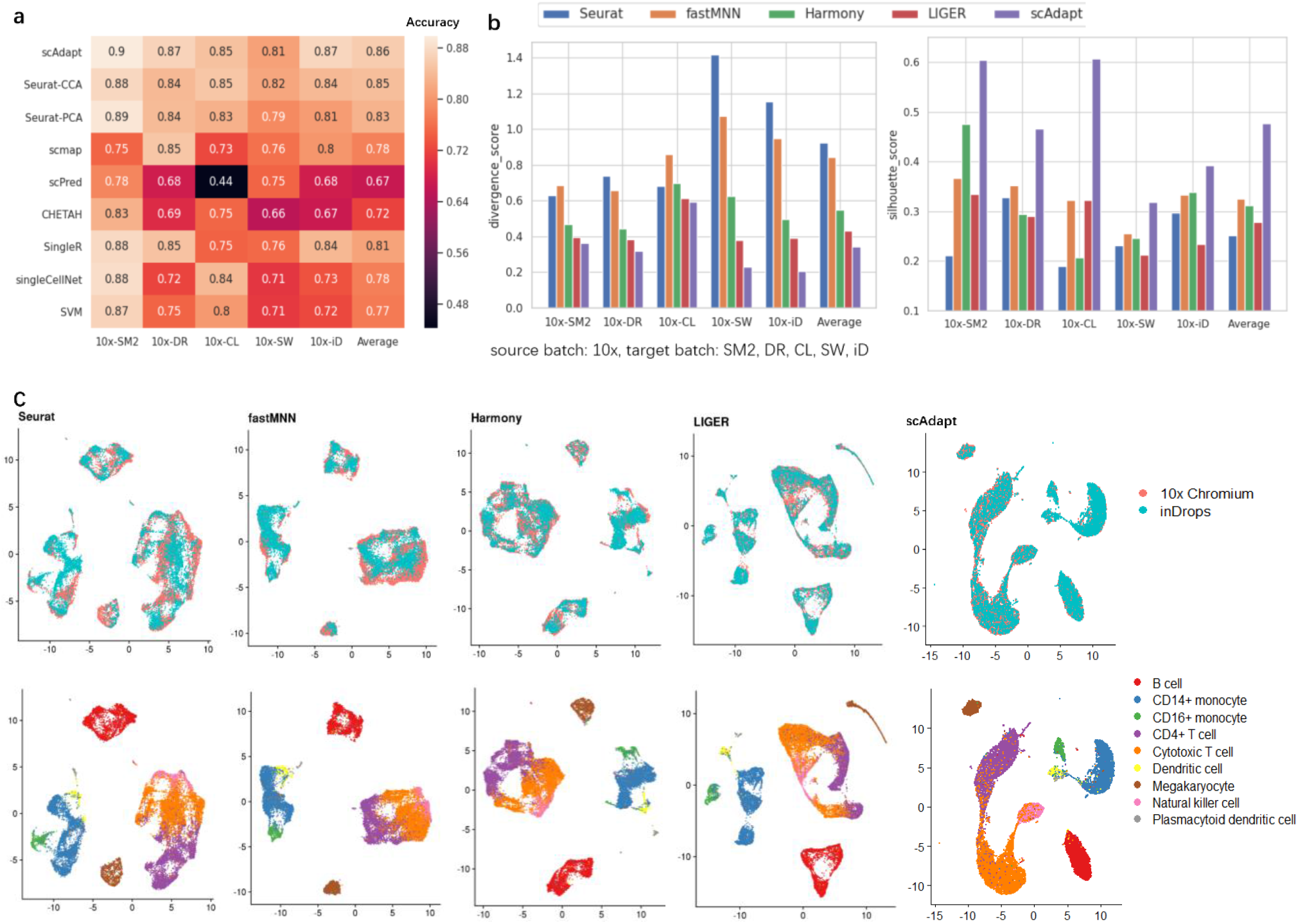
Comparison of classification methods and integration methods for five pairs of cross-platform human PBMC dataset. **a.** Heatmap showing the accuracy of different classification methods. **b.** Divergence score and silhouette score of different integration methods. **c.** UMAP plots of the 10x-iD test pair colored by batch and cell type.

The quantitative performance of the five integration methods are summarized in Fig 3b. As expected, scAdapt is the top method with lowest divergence score and highest silhouette score across all dataset pairs, which is congruent with visualizations plots in Fig 3c, S3. Specifically, divergence score is reduced by 26%-169% on top of Seurat V3, fastMNN, Harmony, and LIGER, respectively. The silhouette score is also improved substantially by 32%-47% when compared to the four competing methods. Although the competing methods can separate the distinct cell types such as B cell and Megakaryocyte well, the highly similar cell types CD4+ T cells, Cytotoxic T cell, and Natural killer cell (Bezman, et al., 2012) are tightly connected in their visualization plots (Fig S3). Since the target data are unlabeled in practice, it would be difficult to visually distinguish them as different cell types. In contrast, scAdapt is able to make the separation of the three clusters visible, which highlights the contribution of the proposed semantic alignment loss and accurate pseudo label.

### 3.3 Performance on cross-species datasets

To demonstrate the ability of scAdapt to more challenging scenario with large distribution difference, we evaluated scAdapt and competing methods by pancreas datasets where the difference come from two sources of variation: species and platform. We used the mouse data from Baron and Tabula Muris cell atlas as source data, and human data from Baron, Segerstolpe, Murano, and Xin as target data. Different from the slight superiority in cross-platform experiment, scAdapt achieves much higher average accuracy (0.93) than the second-ranked scmap (0.81), which confirms the ability of scAdapt to deal with large difference between species. Seurat-CCA, which achieves competitive performance in cross-platform test, suffered an accuracy drop of 22.5% compared to scAdapt. Other six methods are low-performing with accuracy < 0.7. Further inspection of the classification by Sankey plots reveals that the competing methods cannot effectively differentiate some major cell types (Fig S4a). For the alpha cell type, which accounts for the highest proportion (35%) in target data, Seurat-CCA, Seurat-PCA, scPred, and SVM only correctly classified 55%, 26%, 31%, and 65% cells, respectively. For beta cell type with the second highest proportion (26%), 33%-89% cells are correctly categorized by competing methods. By contrast, the accuracy of scAdapt on these two cell types are larger than 0.95. We also performed batch corrected classification evaluations as in cross-platform test and found that the four competing methods can benefit from correction with accuracy of 0.81 (CHETAH), 0.86 (SingleR), 0.78 (singleCellNet), and 0.81 (SVM) (Fig S4b), but are still lower than that of scAdapt (0.93). These results suggest the effectiveness of scAdapt to overcome the variations from both species and platform.

The divergence score and silhouette score computed shows that scAdapt is again the leading method for batch correction (Fig 4b). Similar to simulation and cross-platform scenario, fastMNN ranked second for silhouette score, despite the relatively poor data mixing. Harmony and LIGER produced much lower silhouette scores decreased by 69% and 83%, respectively, compared to scAdapt. Visual inspection reveals that the performance degradation of Harmony and LIGER mainly come from under-correction (Fig 4c, S5). For example, part of the alpha and beta cells are separated as human-specific cell types in xin dataset, which is not consistent with the original assignments. Additionally, scAdapt can clearly separate acinar and ductal cells which come from the same progenitors and are closely associated (Reichert and Rustgi, 2011), while none of the competing methods can separate them in all of the cross-species integration tests. These results suggest that scAdapt is able to maintain biological heterogeneity while effectively reducing unwanted species-specific noise.

**Fig 4.**
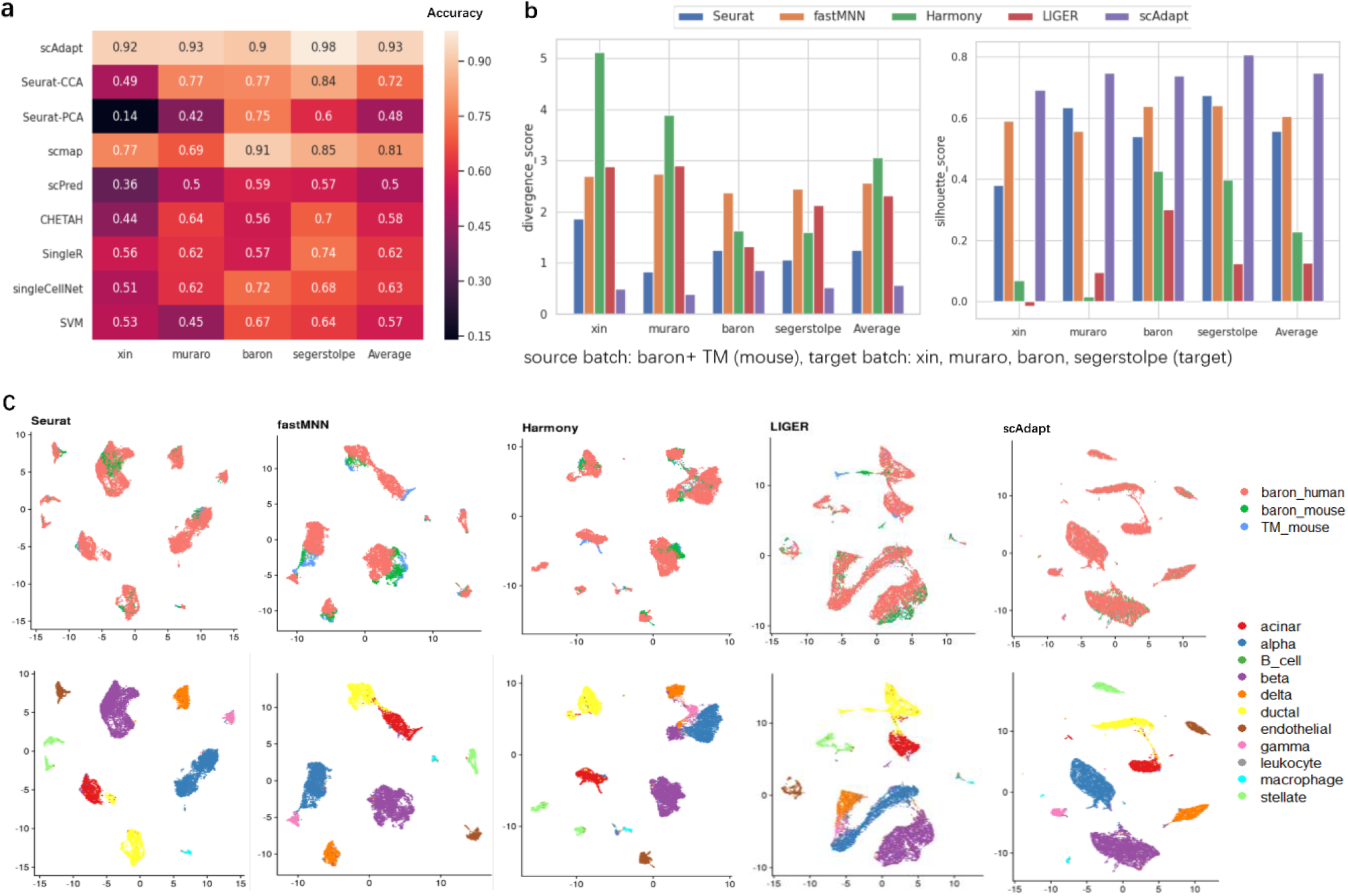
Comparison of classification methods and integration methods for four cross-species pancreas dataset pairs. **a.** Heatmap showing the accuracy of different classification methods. **b.** Divergence score and silhouette score of different integration methods. **c.** UMAP plots of the Baron human dataset mapped to mouse source datasets (Baron and Tabula Muris (TM)) colored by batch and cell type.

Due to the difference between species, the reference data may not contain all cell types in the query data. In order to assess how accurate scAdapt are for discovering new, unknown cell types, we trained it on mouse dataset with “alpha” cells removed, and tested it on human Baron dataset. We labeled the cell “unassigned” if its highest output probability is smaller than 0.5. We found that scAdapt achieved high accuracy for known cell types (accuracy=0.9) while 95% of “alpha” cells are correctly recognized as “unassigned”. Removing “beta” cells obtained similar results with accuracy of 0.95 (known) and 0.89 (“beta”). Pseudolabel thresholding allows new cell types to occupy a distinct region in the embedding. The new cell types in the UAMP visualizations of scAdapt embeddings occupy a distinct region and are clearly distinguishable (Fig S6). These results show that scAdapt is able to identify cell types that are not in the reference dataset.

### 3.4 Application to spatial transcriptomic dataset

Current scRNA-seq technologies require cell dissociation, resulting in losing the spatial localization. Novel spatial transcriptomics methods, such as MERFISH (Moffitt, et al., 2018), can retain cells spatial information, but have a small number of genes that can be simultaneously measured per cell. Thus, finding and merging similar cell types across these two data types is challenging with the limited shared information. To assess how well scAdapt performs compared with alternative methods in this setting, we obtained two datasets profiled from hypothalamic preoptic region of mouse brain, where the dissociated scRNA-seq dataset was sequenced by 10x Chromium as source and the spatial transcriptomics dataset was measured with MERFISH as target. These two datasets have only 154 overlapped genes.

From Fig 5a and S7, we can see that scAdapt achieves an average accuracy of 0.87 over the nine cell types in target dataset, higher than the accuracies of competing classification methods ranging from 0.35 (scPred) to 0.78 (Seurat-CCA). Spatial distribution of the cell types predicted by scAdapt also demonstrate more consistent patterns than competing methods when compared with previous reports (Moffitt, et al., 2018) (Fig S8). For example, the predicted ependymal cells are enriched in a single layer lining the third ventricle. By contrast, this pattern could be missed by Seurat-CCA since it misclassified most ependymal cell as inhibitory neurons. The batch correction results in Fig 5b and S9 suggest that scAdapt successfully mapped cells of the same cell types between the two datasets into a shared embedding, with lower divergence score (0.30) and higher silhouette score (0.47) than alternative approaches (divergence score: 0.33-2.50, silhouette score: 0.20-0.43).

**Fig 5.**
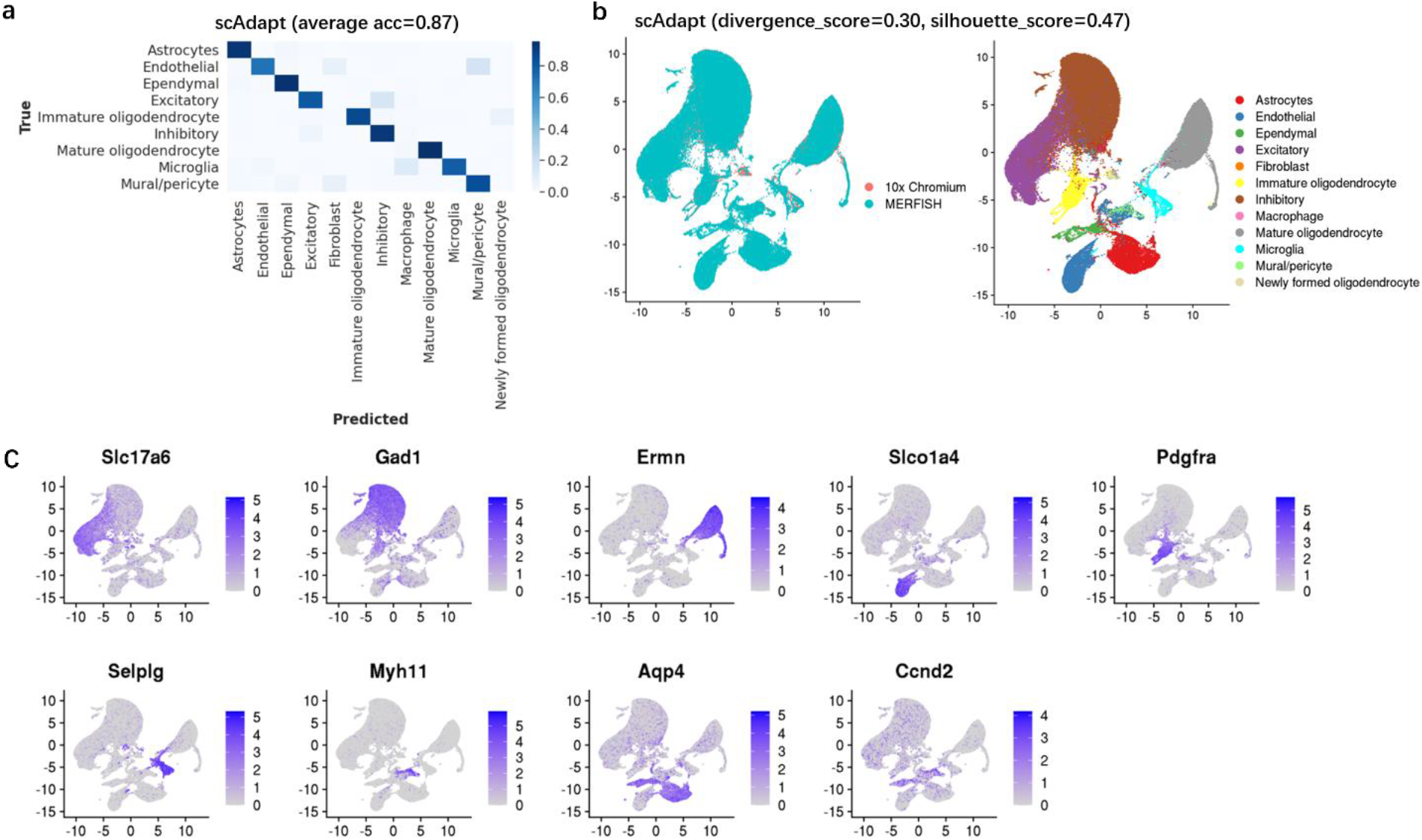
Predicting cell types in spatial transcriptomic dataset (MERFISH) with dissociated scRNA-seq (10x Chromium) dataset from the hypothalamic preoptic region of mouse brain. **a.** Heatmap for the confusion matrix of our method with average accuracy in the bracket. **b.** UMAP plots of the MERFISH dataset mapped to 10x Chromium reference by our method with divergence score and silhouette score in the bracket. Cells are colored by batch (left) and cell type (right). **c.** Expression patterns of the top cell-type-specific marker genes identified by activation maximization in MERFISH dataset. Cells are colored based on the log-normalized expression of marker gene. The gene names are listed in the title of the panel.

To make our neural network model more interpretable, we used activation maximization method to identify the most important gene for predicting certain cell type. We listed the top 10 genes with largest importance score by order in Table S2. We found that among the nine top 1 marker genes identified by our method for each cell type, six genes (Excitatory: *Slc17a6*, Inhibitory: *Gad1*, Immature_oligodendrocyte: *Pdgfra*, Microglia: *Selplg*, Mural_pericyte: *Myh11*, Astrocytes: *Aqp4*) are the same as the ones reported in original publication (Moffitt, et al., 2018), and the other three genes (Mature_oligodendrocyte: *Ermn*, Endothelial: *Slco1a4*, Ependymal: *Ccnd2*) are in maker gene lists of corresponding cell type from PanglaoDB database (Franzén, et al., 2019). In Fig 5c, these genes also exhibited clear expression patterns correlated with corresponding cell types. The results of GO enrichment analysis on the top 10 genes of each cell type were presented in Fig S10. We can see that the selected genes are significantly enriched on GO terms relevant to the biological processes of nervous system, such as glutamate secretion term (GO: 0014047) for excitatory cell, myelination (GO: 0042552) for immature oligodendrocyte cell. These results suggest that the identified genes are consistent with prior biology knowledge and verify the reliability and interpretability of our scAdapt model.

Compared with batch-corrected embedding space, batch correction in the gene expression space is more useful for downstream analysis like the identification of differentially expressed (DE) gene. To address this, we added a decoder layer to reconstruct batch-corrected gene expression from batch-corrected embeddings with mean squared error loss. We used the DE gene detection as a performance measure to evaluate the quality of corrected gene expression. Seurat and fastMNN that produce batch-corrected gene expression matrix were chosen for comparison. We selected DE genes by performing DE analysis between inhibitory cells and all other cells using Wilcoxon rank sum test with logFC> 0.25 and adjust p-value < 0.01. We compared the intersection of DE genes from batch-corrected dataset and uncorrected dataset to evaluate whether the batch correction method can preserve the results of DE analysis on original datasets. From Fig S11, we found that gene expression corrected by scAdapt retained more raw DE genes than those by Seurat and fastMNN (76 vs 67 and 69). Further GO enrichment analysis showed that the DE genes detected by scAdapt were significantly enriched for GO terms relevant to the process of neural communication and development, such as neuropeptide signaling pathway and positive regulation of neuron projection development (Fig S12). These results indicate that scAdapt can benefit from batch-corrected embeddings and accurately retain gene expression features after reconstruction.

## 4 Discussion

In this work, we developed a novel virtual adversarial domain adaptation framework, scAdapt, to perform cell type classification for datasets with batch effects. The virtual adversarial based semi-supervised learning in scAdapt improves classification accuracy using both labeled source dataset and unlabeled target dataset, and domain alignment removes batch effects in the embedding space by making use of label information in the source. For quantitative benchmarks, we used simulated scRNA-seq datasets that vary in the intensity of batch effects, real cross-platform, cross-species, and spatial transcriptomic datasets. Experiments with quantitative measure validated the superiority of scAdapt. Visual inspection also demonstrated that the cells are well mixed and discriminative cluster structure present in the original datasets are preserved. To gain the biological interpretability behind model decisions, we also identified cell type specific marker genes and some of them are validated in PanglaoDB database.

We demonstrated that the comparing classification methods didn’t perform well with large batch difference between source and target dataset. Explicit batch correction can benefit classification in some cases, but the improvement is limited. By contrast, our method is ideal to overcome the strong batch effects according to the experiment results. Additionally, when there are multiple source datasets generated by different sequencing platforms, we can combine these datasets as source, and the cross-platform and cross-species experiment results has shown that our method is robust to the batch effects in the combined dataset. It is to be noted that the source dataset for model training should contain a reasonable number of cells per cell type for reliable cell type annotation. We recommended including at least 10 cells per cell type to adequately represent the transcriptional program as well as variance.

While scAdapt performed best in the experiments, there is still room for improvement. One direction is to enhance the classifier by distinguishing similar subtypes at deeper annotation level since the subtle biological difference between subtypes cannot be recognized easily and often masked by noise from experimental batches and sequencing platforms. Recent advance in fine-grained image classification may serve as a candidate solution which can learn a more discriminative feature representation (Lin, et al., 2015). Another direction is to take the intra-class variance into consideration since the softmax loss tends to collapse all data points from each cell type to a single cluster and thus limits the clustering performance for more detailed subtypes of the labeled cell types. Deep metric learning has shown the ability to maintain the relative relationship of cells (Qian, et al., 2019), and we plan to use this method to preserve local structures for subtype clustering.

