## Supplementary_file for "scAdapt: Virtual adversarial domain adaptation network for single cell RNA-seq data classification across platforms and species"

Table S1. Summary of the dataset pairs used in this study

| Analysis | Source data | | | | Target data | | | |
|---|---|---|---|---|---|---|---|---|
| | # of cells | # of genes | Species | Protocol | # of cells | # of genes | Species | Protocol |
| Splatter simulation | 2000 | 10000 | | | 1000 | 10000 | | |
| Cross-platform PbmcBench | 13028 | 33694 | human | 10x Chromium (v2, v3) | 526 | 33694 | human | Smart-seq2 |
| | | | | | 526 | | | CEL-Seq2 |
| | | | | | 6584 | | | Drop-seq |
| | | | | | 3727 | | | Seq-Well |
| | | | | | 6584 | | | inDrop |
| Cross-species Cross-platform Pancreas | 3429 | 12808 | mouse | inDrop (Baron), SMART-Seq2 (Tabula Muris) | 8506 | 20125 | human | inDrop (baron) |
| | | | | | 2108 | 25525 | | Smart-Seq2 (segerstolpe) |
| | | | | | 2039 | 19127 | | CEL-Seq2 (muraro) |
| | | | | | 1492 | 39851 | | SMARTer (xin) |
| Spatial transcriptomic Mouse brain | 30370 | 22067 | mouse | 10x Chromium | 64373 | 154 | mouse | MERFISH |

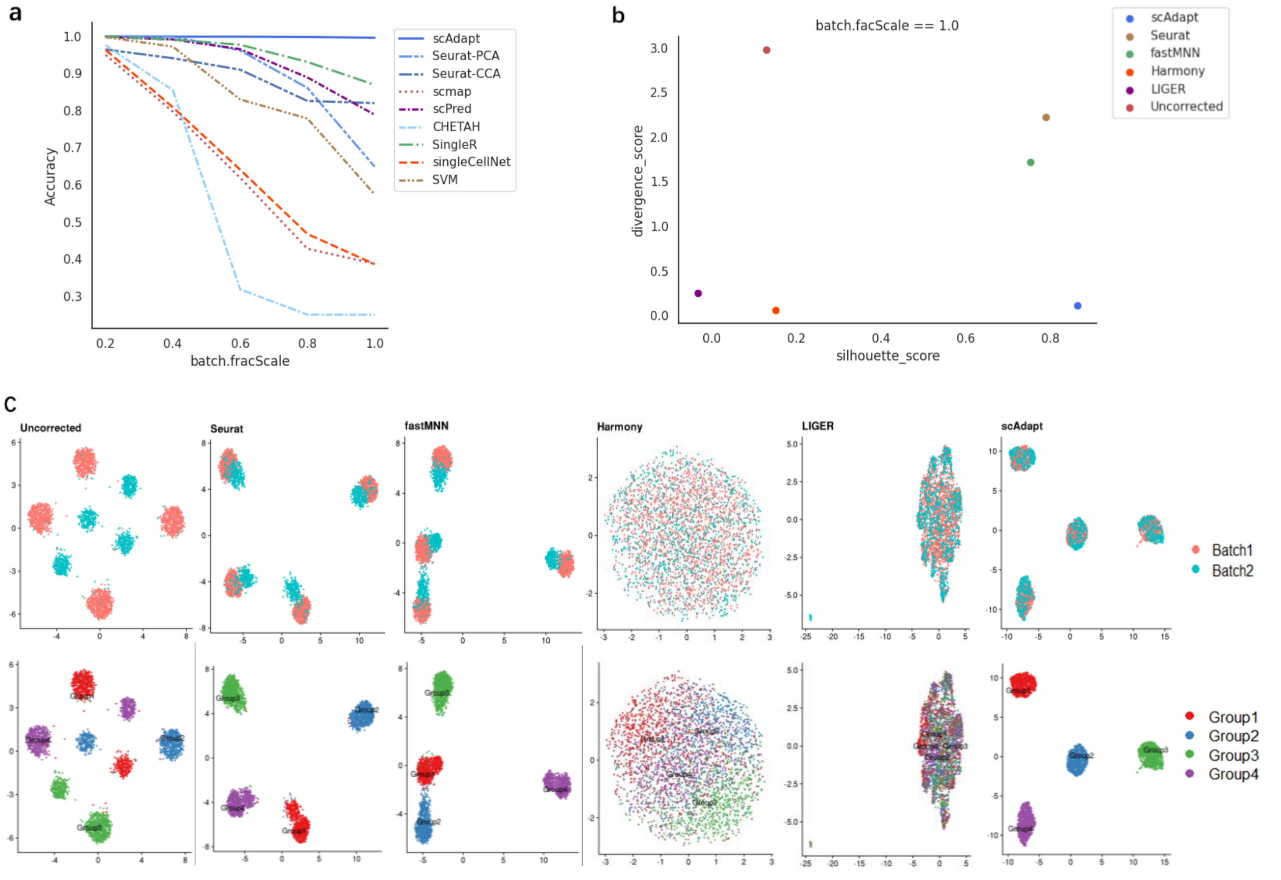


**Fig S1.** Benchmarking of scAdapt against four batch correction methods on simulated data with large batch effects (*batch.fracScale* =1.0) and separable cluster structure (strong clustering signal strength, *de_facScale*= 0.3). **a.** Average accuracy under increasing *batch.fracScale* values where larger values corresponding to stronger batch effects. **b.** The integration quality measured by divergence score versus silhouette score at *batch.fracScale* =1.0. Specifically, a lower divergence score means better cell mixing across datasets and a higher silhouette score indicates better cell type assignment. **c.** UMAP plots colored by batch and cell type at *batch.fracScale* =1.0.

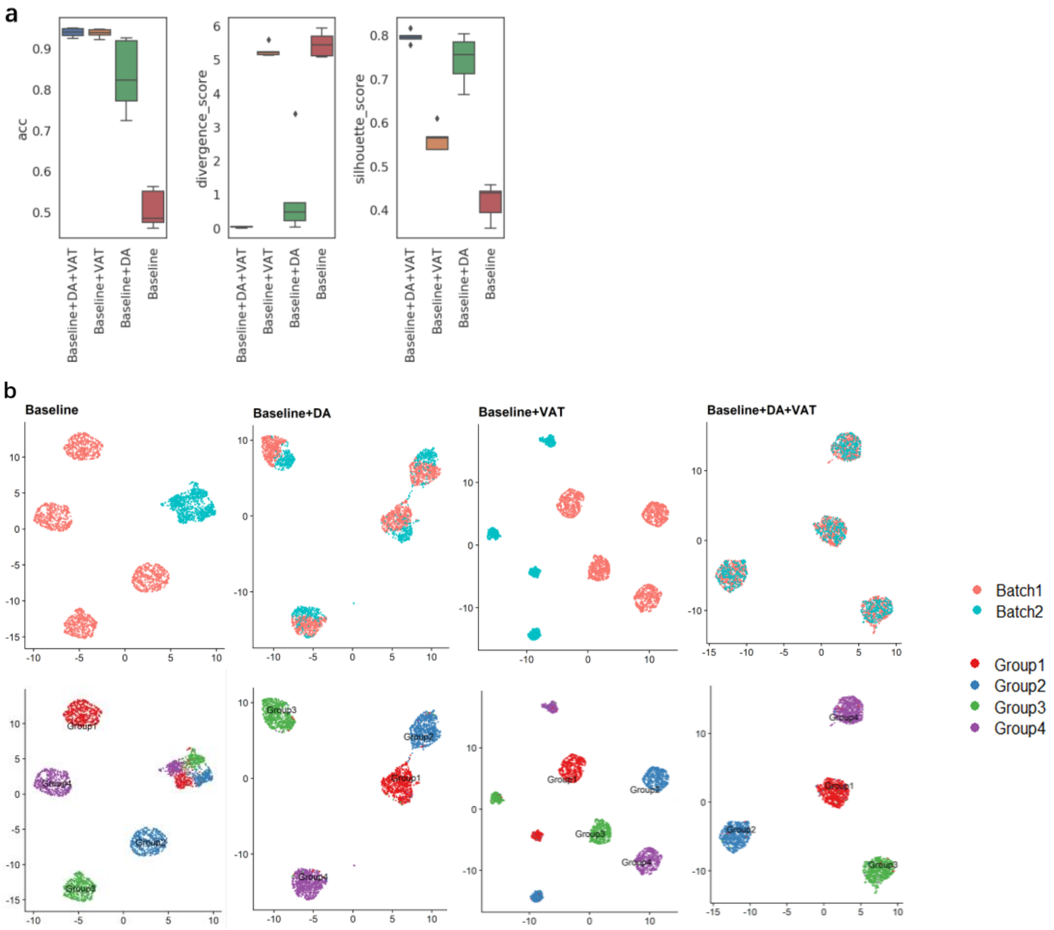


**Fig S2.** Ablation experiments demonstrate that virtual adversarial training (VAT) and global and semantic domain alignment (DA) improve the performance of classification and batch correction, respectively. We evaluated the performance using the most challenging simulation setting for batch correction with batch.fracScale=1.0. **a.** Quantitative evaluation to demonstrate the necessity of different components in scAdapt. Baseline indicates fully-connected neural networks with the same structure as scAdapt but without VAT and DA. **b.** UMAP plots colored by batch and cell type.

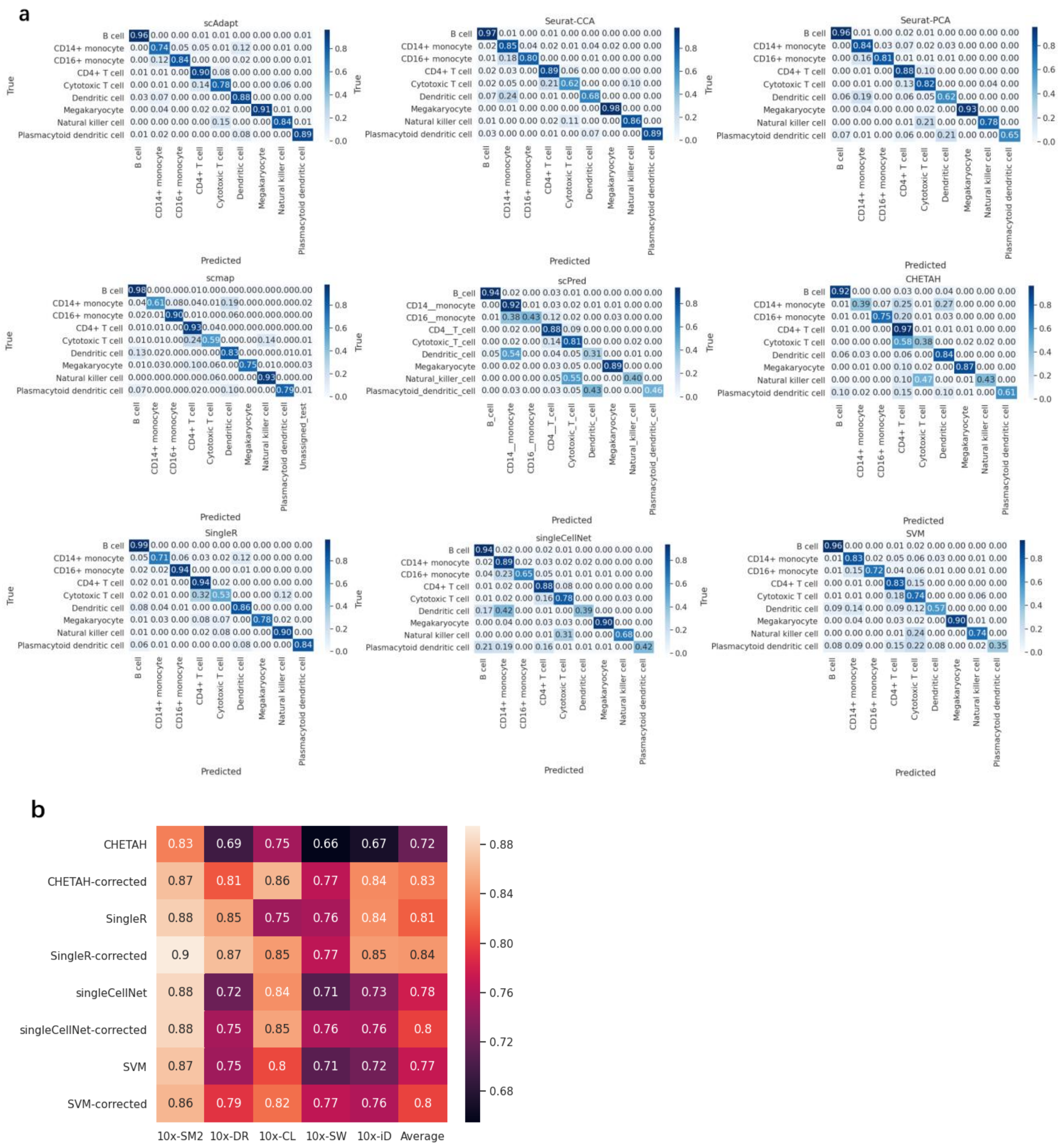


**Fig S2. a.** Heatmap for the overall confusion matrix of the five pairs of cross-platform PBMC datasets. **b.** Heatmap showing the accuracy of CHETAH, SingleR, singleCellNet, and SVM before and after Seurat V3 batch correction. "-corrected" means that the classification method was run over the corrected expression data.

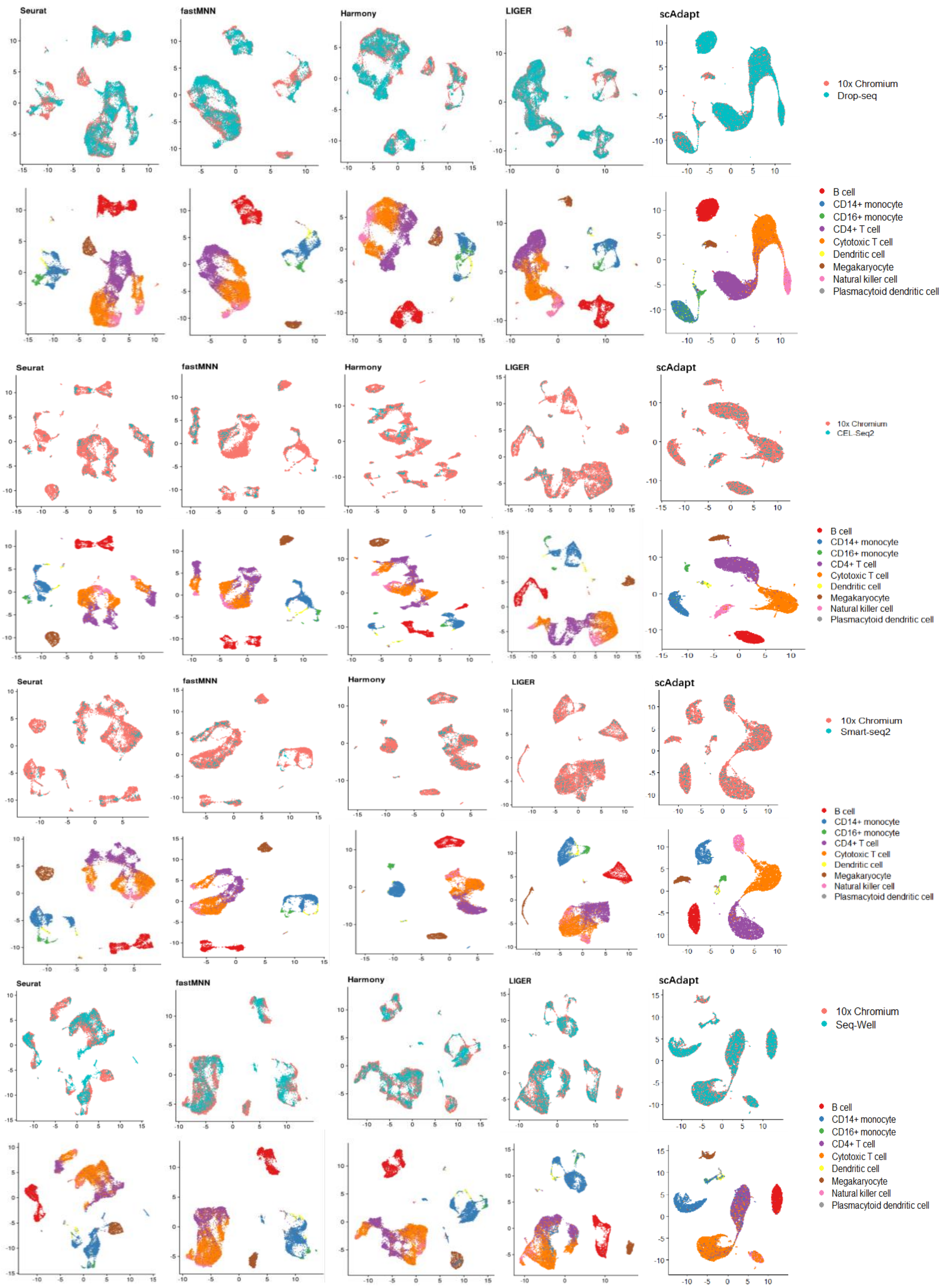


**Fig S3.** UMAP plots of four pairs of cross-platform human PBMC dataset colored by batch and cell type.

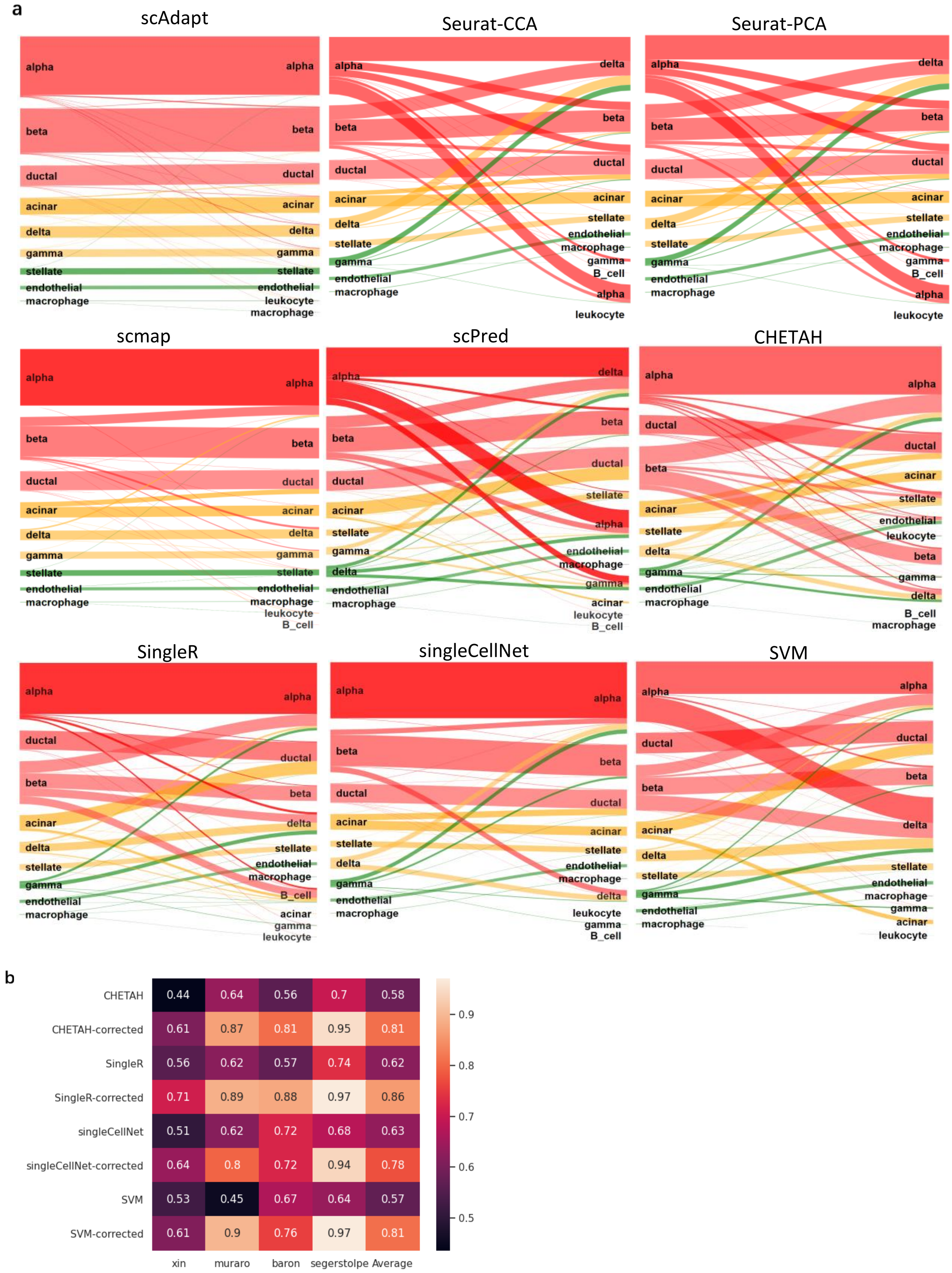


**Fig S4. a.** Sankey plots shows the matching between true labels and predict labels. The overall results of the four pairs of cross-species pancreas datasets were used for plotting. **b.** Heatmap showing the accuracy of CHETAH, SingleR, singleCellNet, and SVM before and after Seurat V3 batch correction. "-corrected" means that the classification method was run over the corrected expression data.

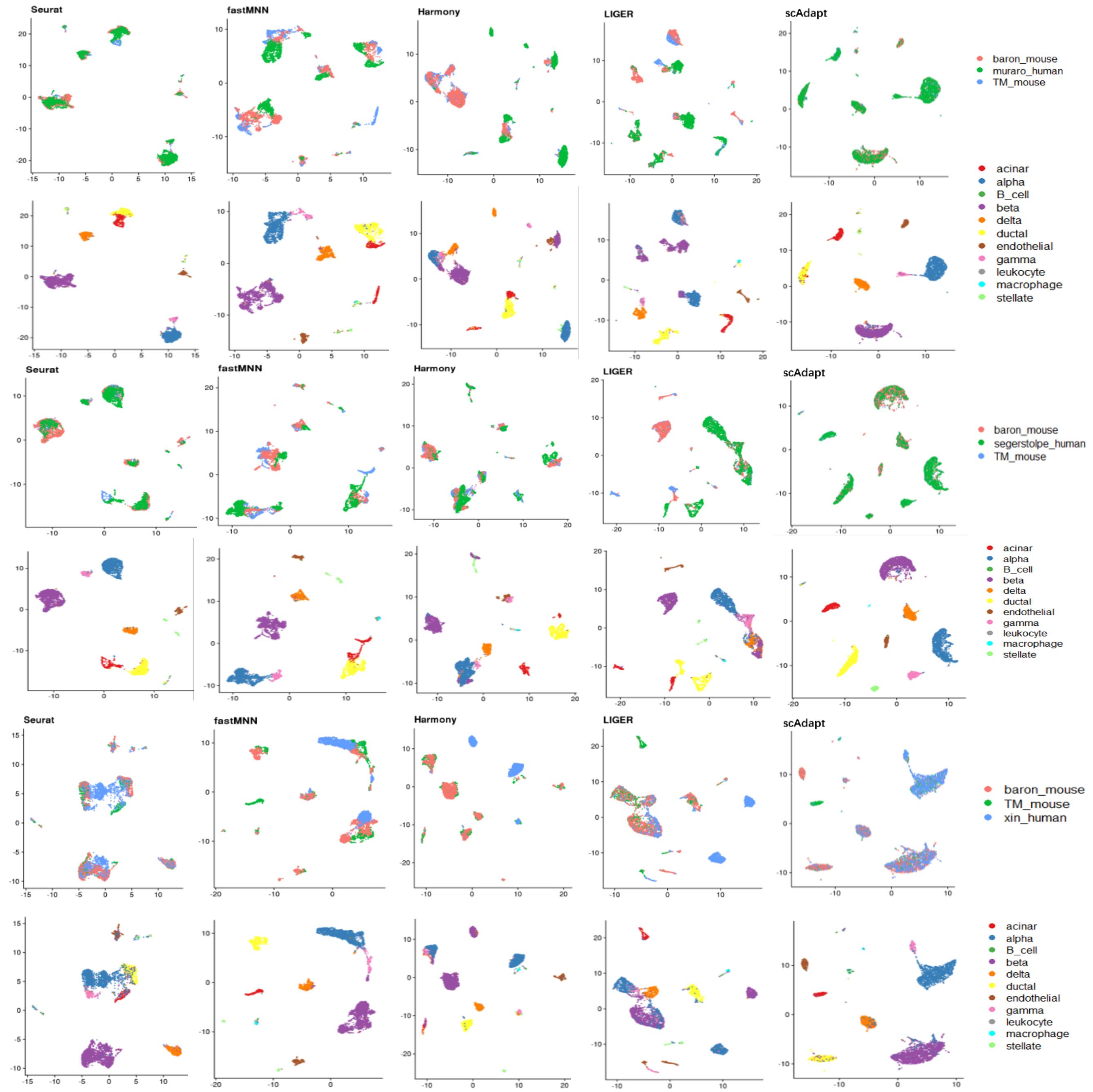


**Fig S5.** UMAP plots of three pairs of cross-species pancreas dataset colored by batch and cell type.

a

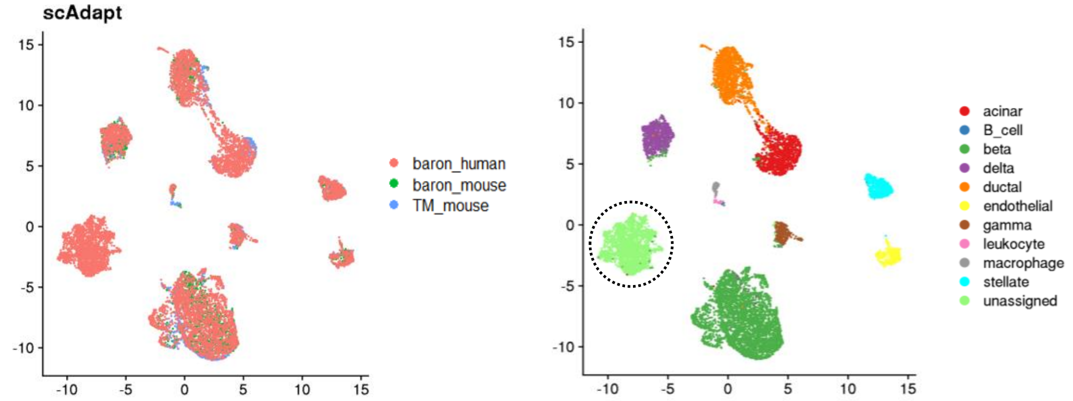


b

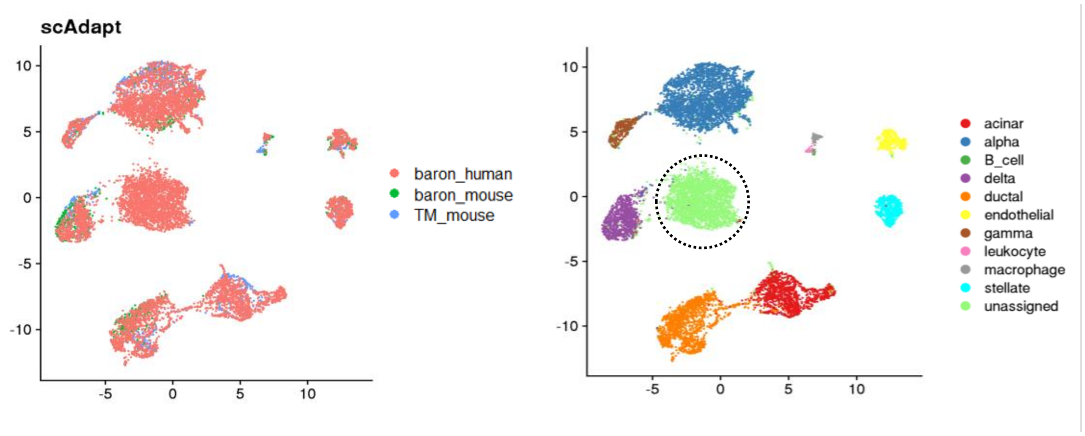


**Fig S6.** Discovering new cell types not in the reference data. **a.** UAMP visualizations of results for reference dataset with "alpha" cells removed. **b.** UAMP visualizations of results for reference dataset with "beta" cells removed. A cell was labelled "unassigned" if its highest probability is smaller than 0.5. "unassigned" cell types are highlight with black circles.

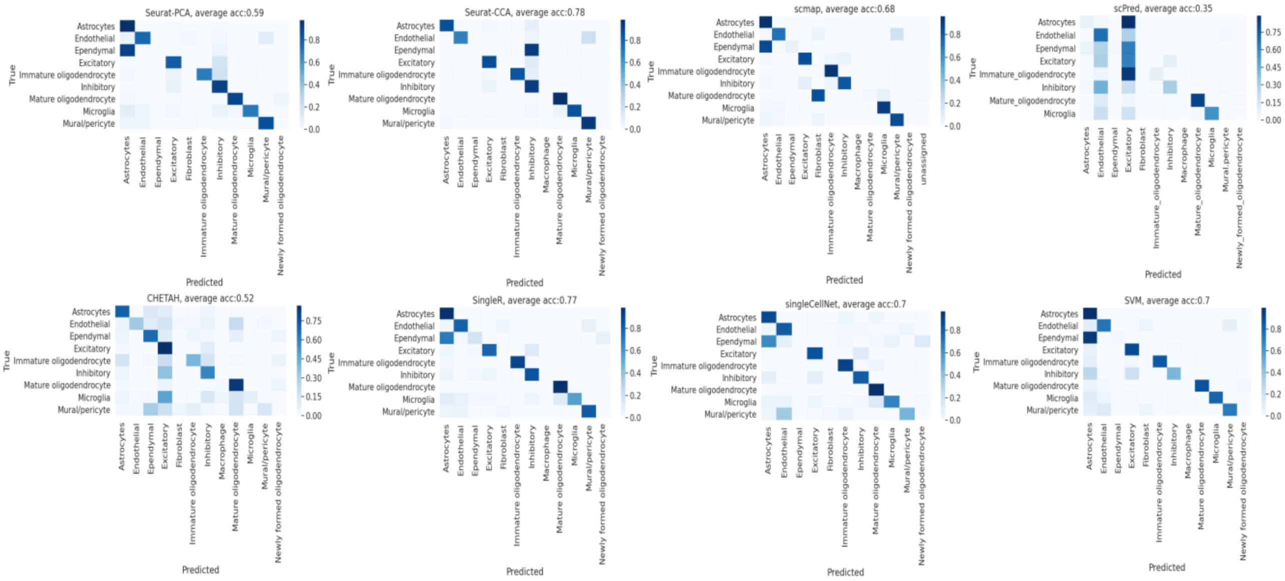


**Fig S7.** Heatmap for the confusion matrix of competing methods on spatial transcriptomic dataset (MERFISH).

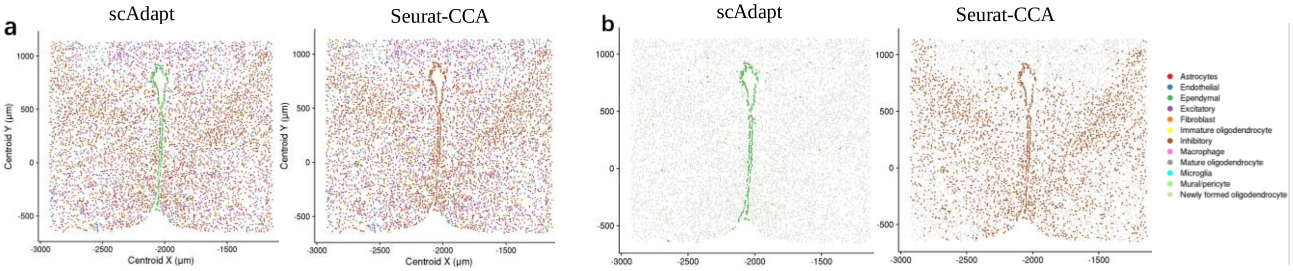


**Fig S8. a.** Spatial locations of the cell types predicted by scAdapt and Seurat-CCA in a 1.8- by 1.8-mm MERFISH imaged slice (Bregma −0.14 mm). Cells are colored by cell types. **b.** Spatial distributions of ependymal cells with distinct patterns are shown as colored dots on the background of other cells shown as gray dots.

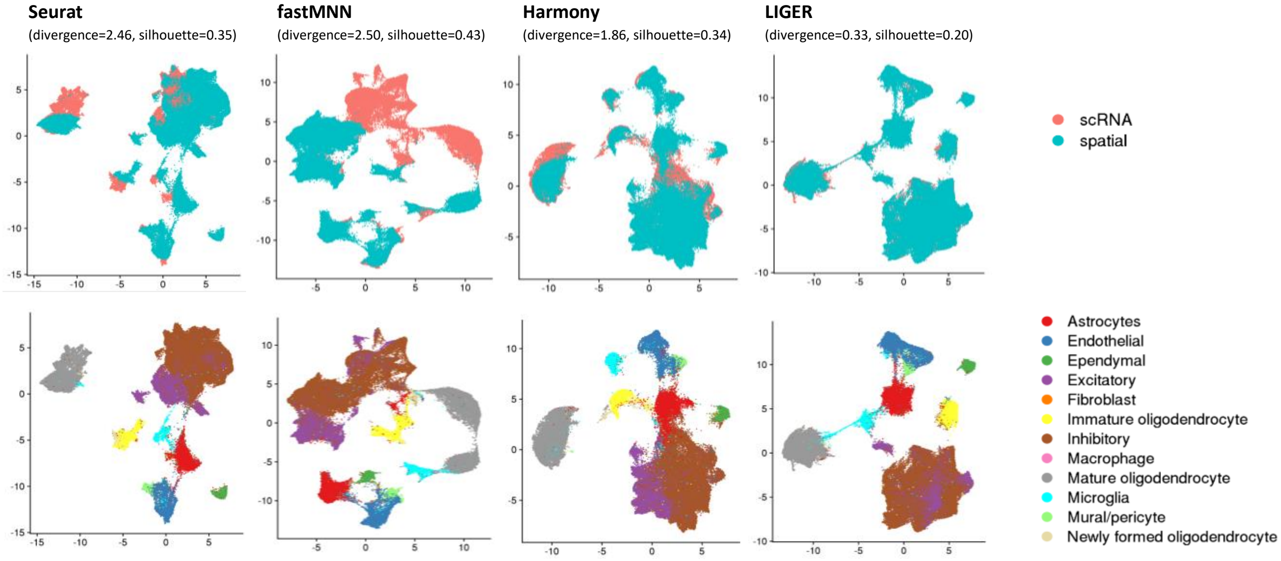


**Fig S9.** UMAP plots of the MERFISH dataset mapped to 10x Chromium dataset by competing methods with divergence score and silhouette score in the bracket. Cells are colored by batch (upper panel) and cell type (lower panel).

| Cell types | Top 10 genes identified by activation maximization |
|---|---|
| Excitatory | *Slc17a6 Scg2 Mbp Syt4 Pak3 Gabra1 Gad1 Trh Nnat Bdnf* |
| Inhibitory | *Gad1 Mbp Scg2 Syt4 Pak3 Cpne5 Gabra1 Penk Baiap2 Nnat* |
| Mature_oligodendrocyte | *Ermn Mbp Etv1 Opalin Gjc3 Sgk1 Ttyh2 Ndrg1 Lpar1 Plin3* |
| Endothelial | *Slco1a4 Mbp Cdkn1a Klf4 Scg2 Sema3c Ndrg1 Cyr61 Opalin Ccnd2* |
| Immature_oligodendrocyte | *Pdgfra Cspg5 Mbp Gjc3 Nnat Sox4 Cyr61 Traf4 Baiap2 Scg2* |
| Microglia | *Selplg Slc15a3 Mbp Sgk1 Cdkn1a Rgs2 Man1a Scg2 Nnat Sst* |
| Mural_pericyte | *Myh11 Rgs5 Mbp Ccnd2 Scg2 Lmod1 Nnat Sox8 Slco1a4 Cyr61* |
| Astrocytes | *Aqp4 Mbp Cxcl14 Nnat Mlc1 Cspg5 Gabrg1 Tiparp Pou3f2 Sox8* |
| Ependymal | *Ccnd2 Nnat Mlc1 Mbp Cspg5 Cyr61 Tiparp Gabrg1 Plin3 Cbln2* |

**Table S2.** Top 10 genes identified by activation maximization for scAdapt model on MERFISH-10x dataset.

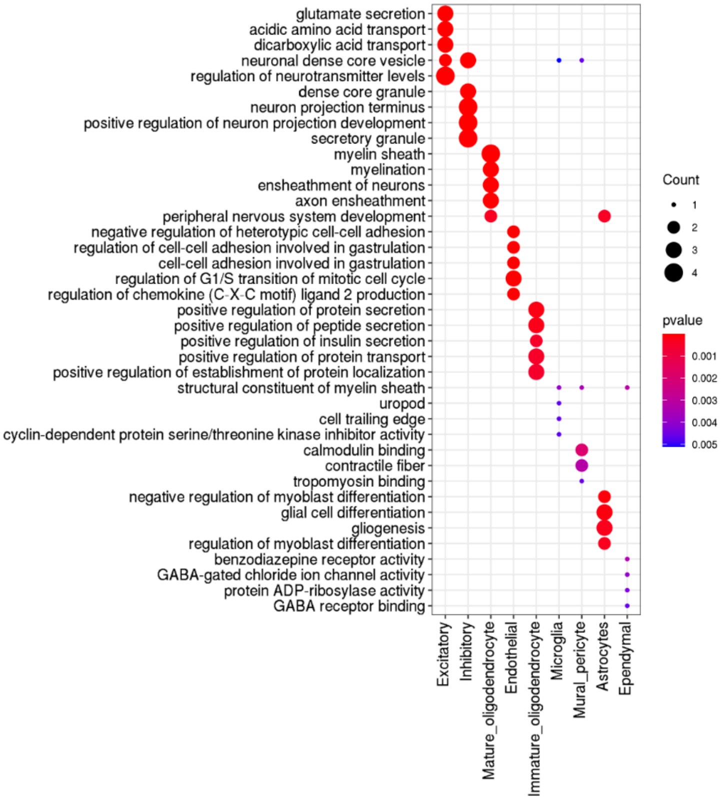


**Fig S10.** Dot plots of the top 5 most enriched Gene Ontology (GO) terms from cell-type specific GO enrichment analysis with the top 10 marker genes identified by activation maximization method in Table S2.

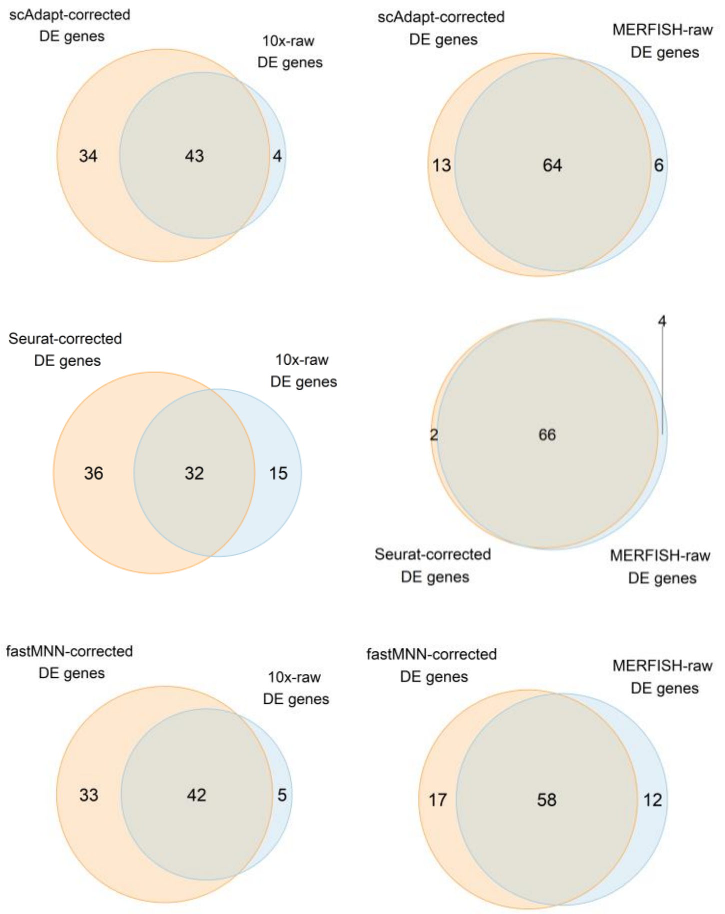


**Fig S11.** Venn diagrams representing the intersection of differential expression (DE) genes identified in batch-corrected (scAdapt, Seurat, and fastMNN) (orange circle) and raw MERFISH-10x dataset (blue circle). DE genes are selected by performing DE analysis between inhibitory cells and all other cells using Wilcoxon rank sum test with logFC > 0.25 and adjust p-value < 0.01.

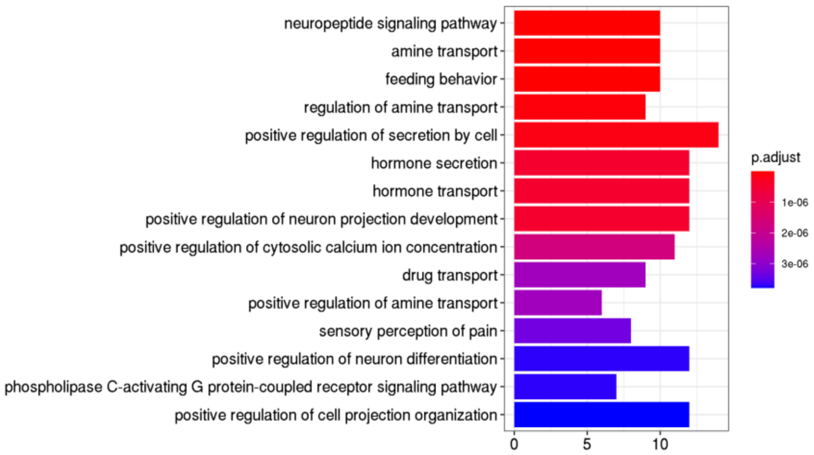


**Fig S12.** GO-enrichment analysis of differential expression genes identified from MERFISH-10x dataset corrected by scAdapt. DE genes are selected by performing DE analysis between inhibitory cells and all other cells using Wilcoxon rank sum test with logFC > 0.25 and adjust p-value < 0.01.
